# DNA-PAINT single-particle tracking (DNA-PAINT-SPT) enables extended single-molecule studies of membrane protein interactions

**DOI:** 10.1101/2022.08.25.503948

**Authors:** Christian Niederauer, Chikim Nguyen, Miles Wang-Henderson, Johannes Stein, Sebastian Strauss, Alex Cumberworth, Florian Stehr, Ralf Jungmann, Petra Schwille, Kristina A. Ganzinger

## Abstract

DNA-PAINT based single-particle tracking (DNA-PAINT-SPT) has recently significantly enhanced observation times in *in vitro* SPT experiments by overcoming the constraints of fluorophore photobleaching. However, with the reported implementation, only a single target can be imaged and the technique cannot be applied straight to live cell imaging. Here we report on leveraging this technique from a proof-of-principle implementation to a useful tool for the SPT community by introducing simultaneous live cell dual-colour DNA-PAINT-SPT for quantifying protein dimerisation and tracking proteins in living cell membranes, demonstrating its improved performance over single-dye SPT.

---

Single-particle tracking (SPT) is a powerful method to investigate the orchestration of biomolecular processes at cell membranes or in reconstituted systems ^1,2^. To detect and follow the molecules of interest, these are usually fluorescently labelled, with observation times and localisation precision depending on label brightness and photostability. Furthermore, labels have to be conjugated with the target molecule in 1-1 stoichiometry to obtain meaningful data ^3^. Due to their small size, brightness and ease of chemical addressability, organic dyes conjugated to genetically encoded protein tags are currently the preferred labelling strategy for SPT ^4,5,6,7^. However, observations are typically only possible for a few seconds at 20–50 nm spatial precision before the dyes photobleach ^2^. Short trajectories particularly hamper quantitative studies of molecular association by multi-colour SPT, as they reduce the dynamic range of these experiments and make it hard to distinguish true co-diffusion events from chance encounters.

Recently, we have demonstrated how DNA-PAINT-SPT can increase trajectory lengths by circumventing the limited photon budget of single dyes ^8^. In DNA-PAINT-SPT, short dye-labelled DNA oligonucleotides (imager strands) transiently bind to a target-bound complementary docking strand that contains several repeating with speed-optimised sequences ^9,10^. As multiple imager strands can thus bind simultaneously and are designed to exchange on a time scale similar to that of dye photobleaching, this allowed us to follow the motion of DNA-origami on a supported lipid bilayer (SLB) for minutes rather than seconds ^8^. While the concept of constantly exchanging fluorophores to prevent photobleaching has gained traction in the field of single-molecule fluorescence ^11,12^, it is yet to be implemented in more complex biological samples.

Here, we introduce a new motif for dual-colour DNA-PAINT-SPT for measuring protein-protein interactions at the single-molecule level, and use it to reliably quantify ligand-induced protein dimerisation in membranes. We further extend our new dual-colour DNA-PAINT-SPT implementation to live cell imaging applications and demonstrate its improved performance over single-dye SPT.

To apply DNA-PAINT-SPT in dual-colour experiments of molecular interactions, we designed orthogonal docking-imager strand pairs (Fig. 1a, Supplementary Note A) that exhibited negligible crosstalk (Fig. 1b). We then used our dual-colour DNA-PAINT-SPT approach to study the homodimerisation of the FK506 binding protein (FKBP) reconstituted on SLBs in the presence of the dimerisation agent AP20187^13,14^. We reconstituted His-tagged FKBP-SNAPtag fusion proteins on SLBs containing nickel-nitrilotriacetic acid (Ni-NTA) lipids, and labelled the proteins with orthogonal benzylguanine (BG)-modified DNA docking strands via their SNAPtag. When we added the respective imager strands in solution (Fig. 1c), we reliably detected co-diffusion events and hence protein dimerisation (Fig. 1d-e, Fig. SV1, Fig. SV2) using a total internal reflection fluorescence microscope (TIRFM). Individual dimers could routinely be followed for tens of seconds, often even for several minutes (Fig. 1f-g; Fig. S1, Fig. SV3), in stark contrast to various single-dye labelled controls, including state-of-the-art fluorophores developed for single-molecule imaging ^15^ (Fig. 1f-g; Fig. S2). Similarly, the number of observable DNA-PAINT labelled proteins remained stable on the time scale of video acquisition (*≈* 6 min) in contrast to single-dye controls: not only does the number of detected dimers rapidly decrease over time for single-dye labelling, but also the apparent dimer lifetime is shortened as a consequence, resulting in a systematic underestimation of dimer stability (medians of co-diffusion durations measured with DNA-PAINT and single-dye labelling: *T*_PAINT_ = 24.2 ± 3.8 s, and *T*_SD_ = 4.1 *±* 1.3 s; Fig. 1g). As a result of the frictional drag incurred by the increased membrane footprint of the dimers, we observed slowing down of diffusion of dimerised proteins by 28 % (Fig. 1h). Titrating the ligand concentration, we were able to extract 2D dissociation constants *K_B_* (ligand from solution binding to FKBP monomer; [M]) and *K_X_* (cross-linking of a ligand-bound monomer with a free monomer; [µm^−2^]), obtaining an excellent fit for an analytical homodimerisation model to our DNA-PAINT single-molecule data (*K_B_* = 0.85 *±* 0.17 nM, *K_X_* = 2.6 *±* 0.2 × 10^−2^ µm^−2^; Fig. 1i, Supplementary Note B, ^16^). Inducing dimerisation with an anti-SNAPtag antibody instead of the AP20187 ligand was reflected in a drastic slow down of diffusion upon dimerisation (two-fold reduction compared to AP20187-induced dimers, see Fig. S3, Fig. S4) and larger dissociation constants (*K_B_* = 136 *±* 41 nM, *K_X_* = 0.12 *±* 0.02 µm^−2^; Fig. S5, Fig. S6), in line with the expectations from solution kinetics predicting weaker affinities for the antibody.

**Figure 1.**
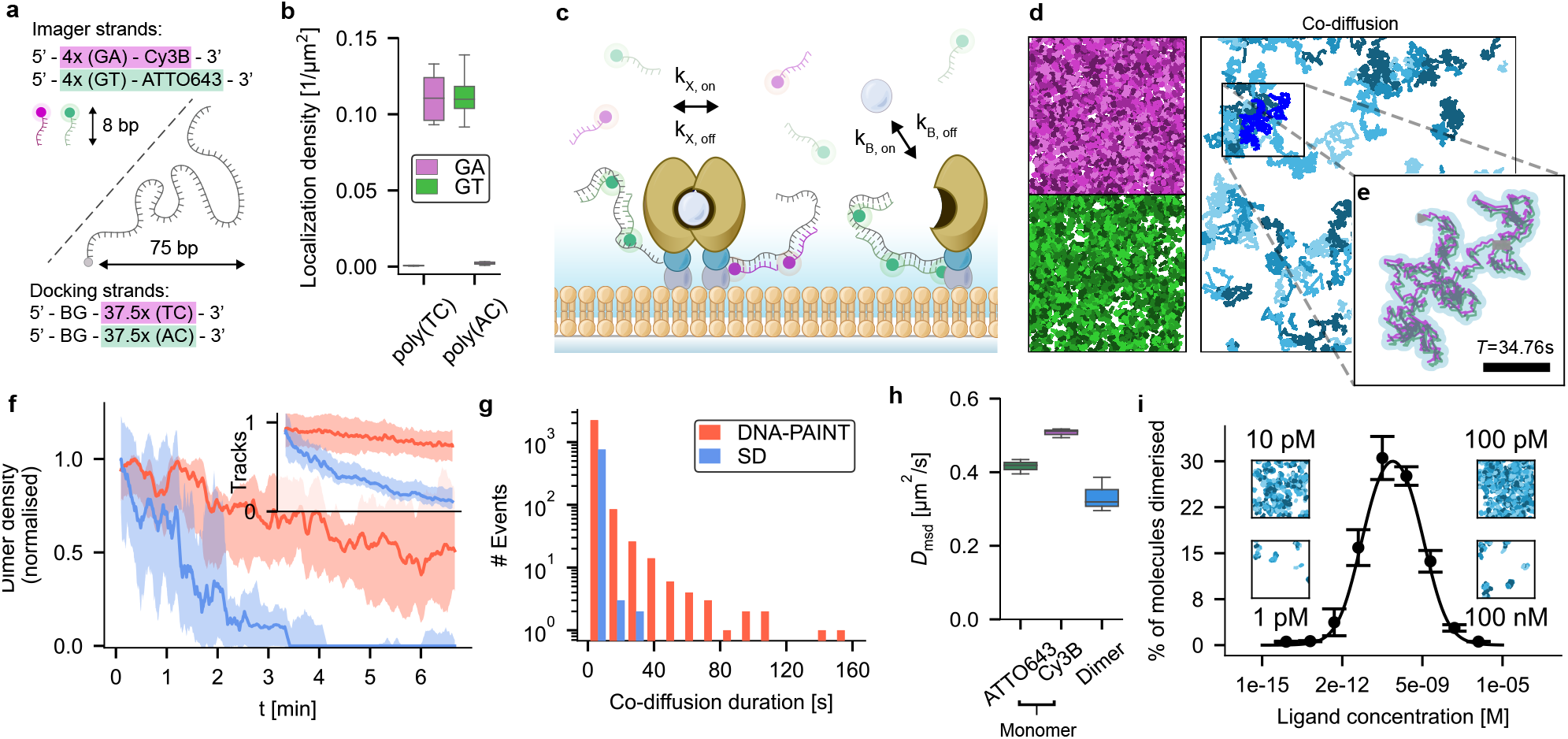
DNA-PAINT-SPT allows quantitative single-molecule studies of FKBP homodimerisation. a) Sequence design for dual-colour DNA-PAINT-SPT. Imager strands are short fluorescently labelled single-stranded DNA consisting of nucleotide-pair repeats (GA and GT). Docking strands are 75 nucleotide long single-stranded DNA, consisting of repeats of the respective complementary nucleotide-pairs (TC and AC) and a benzylguanine (BG) moiety for covalent labelling of proteins with a SNAPtag. A single docking strand can be occupied by several imager strands at the same time, with binding times tuned to match bleaching kinetics, allowing for the continuous observation of labelled proteins. b) Density of tracks of reconstituted SNAPtag fusion proteins labelled with BG-conjugated orthogonal docking strand sequences and their respective imager strands conjugated to ATTO643 or Cy3B fluorophores. Boxes, line and whiskers show, respectively, 25–75 quartiles, median, and minimum and maximum values of data pooled from three fields of view of duplicate samples per condition. c) Schematic of the *in vitro* dimerisation assay. FKBP-proteins (yellow) are reconstituted on a supported lipid bilayer and labelled via their SNAPtag (blue) with orthogonal DNA oligonucleotide docking strands. Short complementary imager strands, conjugated to Cy3B (magenta) and ATTO643 (green) fluorophores transiently bind to the docking strand and allow for dual-colour single-particle tracking. Dimerisation of FKBP-proteins is induced by adding a dimerisation agent (grey sphere). d) Single-molecule trajectories collected during a 40 seconds recording (left, magenta and green) and detected dimerisation events (right, blue). Fields of view are 70 µm *×* 70 µm. e) Tracks of two monomers (magenta and green) co-diffusing for 34.76 seconds (869 frames). For displaying purposes, tracks were moved in opposite *x* and *y* directions by 0.2 µm. Co-diffusion as detected by the tracking algorithm is presented as a blue-shaded track. Scale bar: 5 µm. f) Mean number of dimers per frame observable using DNA-PAINT-SPT (red) or single-dye control (blue), normalised to their respective initial values. Inset: Number of tracks using DNA-PAINT-SPT (red) or single-dye control (blue), normalised to their respective initial values. Curves represent the median with shaded areas indicating the 25-75 quartiles of data collected from three samples per condition. g) Histogram of interaction durations for ligand-induced dimerisation as detected during six minute measurements, using DNA-PAINT (red, mean = 24.2 *±* 3.8 s) or single-dye labelling (blue, mean = 3.3 *±* 1.3 s). Data collected during measurements of six minutes each from three samples per condition. h) Diffusion constants of monomers and dimers labelled with DNA-PAINT or single dyes. Boxes, line and whiskers show, respectively, 25–75 quartiles, median, and minimum and maximum values of data collected from three samples per condition. i) Fraction of dimerised molecules as detected using DNA-PAINT-SPT during ligand titration experiments. Fitting the data results in dissociation constants *K*_*X*_ = 5.9 *±* 0.5 *×* 10^−3^ µm^−2^, *K*_*B*_ = 33 *±* 5 pM. Insets: Trajectories of detected dimers for selected ligand concentrations, collected during 40 seconds measurements. Fields of view are 70 µm *×* 70 µm. Error bars denote mean *±* standard deviation of data collected on five fields of view per condition, samples were prepared independently as duplicates.

Having developed dual-colour DNA-PAINT-SPT, we next sought to establish DNA-PAINT-SPT for live cell SPT experiments. Since DNA-PAINT-SPT relies on the diffusive exchange of fluorescent imager strands from solution, a surface-restricted excitation geometry (i.e. TIRFM) is required to suppress the background signal from free imager strands. However, the implementation of TIRFM for DNA-PAINT-SPT on live cells is not trivial: the surface properties need to be tuned to facilitate cell adhesion while allowing imagers to diffuse underneath the cell (Fig. 2a). At the same time, unspecific binding of free imagers has to be minimal. Out of all passivation methods screened, we found that SLBs containing lipids modified with a integrin-recognition peptide (DSPE-PEG2000-RGD) suppressed nonspecific binding the best while promoting cell attachment and imager strand diffusion underneath the cells (Fig. 2a and Supplementary Note C, Fig. S7, Fig. SV5). For selected fluorophores, passivation using a non-covalent PLL-PEG/PLL-PEG-RGD coating was also sufficient (Supplementary Note C). After surface optimisation, we labelled a model transmembrane protein (SNAP-CD86tm-FKBP-GFP) via its extracellular SNAPtag with a BG-DNA docking strand for DNA-PAINT-SPT (Fig. 2b): after addition of complementary imager strands carrying Cy3B fluorophores, individual membrane proteins on the cell surface are visible as diffusing bright fluorescent spots with step-wise fluctuating intensity, as expected from the continuous binding and unbinding of imager strands (Fig. 2b-d; Fig. SV4). We note cells appear as dark shadows surrounded by elevated background (Fig. 2c) in our TIRFM videos, indicating that diffusion of imager strands into the space between cells and glass coverslips is restricted. However, using a 75 nucleotides DNA docking strand and 40 nM of imager strands, we could achieve on-rates sufficient for continuous exchange. Comparing DNA-PAINT-SPT on these cells to single-dye controls, we found that the number of observable DNA-PAINT labelled membrane proteins remained stable (*>* 85% after six minutes) while for the single-dye control this number decreased to less than a fifth of the initial value (*<* 15% after six minutes), with most of the remaining observable molecules diffusing in from the cell boundaries (Fig. 2e-f, Fig. S8). As a combined measure for average trajectory length and number, we plot the number of tracks that are longer than a given threshold time *T*, normalised to the number of molecules initially detected per cell, for DNA-PAINT and single-dye labelled membrane proteins (Fig. 2g, Fig. S9). This shows that also for live cell imaging, DNA-PAINT-SPT not only keeps the number of observable molecules constant for long durations (> 6 min), but also increases the duration of individual trajectories (DNA-PAINT: *τ*_1*/*2_ = 31 *±* 13 s; single-dye: *τ*_1*/*2_ = 17 *±* 3 s). The diffusion constants for both DNA-

**Figure 2.**
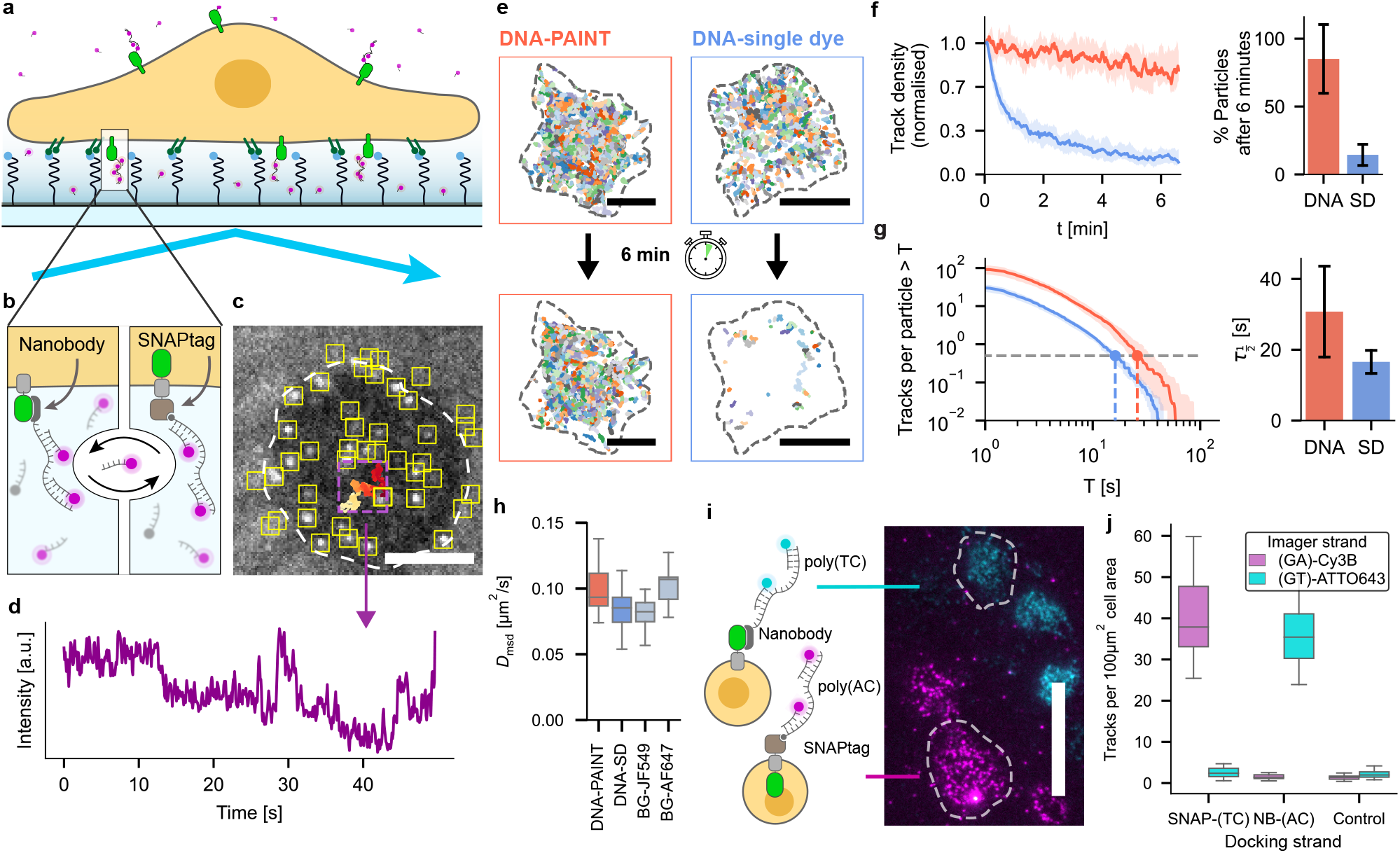
DNA-PAINT-SPT enables extended tracking of individual membrane proteins on live cells. **a)** Schematic of DNA-PAINT-SPT experiment: cells rest on PEG-cushioned SLBs decorated with RGD peptides for optimal cell attachment, imager access and surface passivation. **b**) Membrane proteins of interest are labelled with a oligonucleotide (docking strand) using protein tags or nanobodies, allowing continuous binding and unbinding of several fluorescently-labelled, complementary imager strands from solution. **c**) Video frame of TIRFM video showing single-molecule trajectories of DNA-PAINT labelled membrane proteins expressed on a Jurkat T cell. Localised molecules (yellow boxes) and trajectory of an individual molecule (magenta box, trajectory is colour-coded in time from red to yellow). Scale bar: 5 µm. **d**) Intensity of single molecule trajectory displayed in Fig. 2c, magenta box) over time. **e**) Membrane protein (SNAPtag-CD86tm-FKBP-GFP) trajectories collected during the first (top) and last (bottom) 20 seconds of a six minute recording; Left: membrane proteins labelled with DNA-PAINT (BG-DNA, nine repeats of eight nucleotides complementary to Cy3B-labelled imager strand). Right: membrane proteins labelled with DNA single-dye control (BG-DNA, 35 nucleotides complementary to Cy3B-labelled imager strand). Scale bar: 10 µm. **f**) Number of trajectories per frame on individual cells labelled with DNA-PAINT (red, *n*_cells_ = 25) or single dyes (blue, *n*_cells_ = 32), normalised to initial number of trajectories. **g**) Single-molecule trajectories on individual cells labelled with DNA-PAINT (red, *n*_cells_ = 25) or single dyes (blue, *n*_cells_ = 32) with a duration longer than *T*, normalised to the initial number of trajectories per cells. In **f** and **g**, curves represent the median with shaded areas indicating the 25-75 quartiles of data collected from three samples per condition. Bars and error bars denote mean and standard deviation. **h**) Diffusion constants of cell membrane proteins labelled with DNA-PAINT (*n*_cells_ = 28), DNA single-dye control (*n*_cells_ = 43) and organic fluorophores JF549 (*n*_cells_ = 21) and AF647 (*n*_cells_ = 7). Boxes, line and whiskers show, respectively, 25–75 quartiles, median, and minimum and maximum values of data. **i**) Cells expressing either GFP-CD86tm-FKBP-HALOtag (cyan) or SNAPtag-CD86tm-FKBP-GFP (magenta) fusion proteins labelled with GFP-nanobody-conjugated docking strands (nanobody-DNA, nine repeats of eight nucleotides complementary to Cy3B-labelled imager strand) or with BG-DNA (BG-DNA, nine repeats of eight nucleotides complementary to ATTO643-labelled imager strand). Scale bar: 20 µm. **j**) Density of tracks detected on cells expressing extracelullar SNAPtag-(*n*_cells_ = 38) or GFP-fusion proteins (*n*_cells_ = 35) and non-transfected controls (*n*_cells_ = 30), when labelled in parallel with BG-or nanobody-conjugated docking strands with orthogonal sequences and their respective imager strands. Boxes, line and whiskers show, respectively, 25–75 quartiles, median, and minimum and maximum values of data.

PAINT and single-dye DNA labelled proteins were similar to the direct labelling of the SNAPtag with BG-conjugated organic fluorophores (DNA-PAINT with Cy3B-labelled imager strands: 0.093 ± 0.017 µm^2^/s, single-dye DNA labelled with Cy3B: 0.085 ± 0.014 µm/s, BG-JF549: 0.082 ± 0.020 µm/s, BG-AF647: 0.107 ± 0.016 µm/s; Fig. 2h, Fig. S10). This suggests that DNA-PAINT-labelling *per se* influences diffusion to a lesser extent than the choice of fluorophore ^17^. We also note that we did not observe any exclusion effects of DNA-labelled proteins from cell-surface contacts (Fig. 2c, **??**). Notably, DNA-PAINT-SPT worked equally well with a second, non-covalent labelling approach, using a docking strand conjugated to an antiGFP-nanobody (Fig. 2b,i-j). Thus, we can use two orthogonal labelling approaches (BG-conjugated docking strands or antiGFP-nanobodies) in combination with the orthogonal docking-imager pairs for dual-colour DNA-PAINT-SPT on live cells (Fig. 2i-j; Fig. SV6).

In conclusion, we present DNA-PAINT-SPT as a promising technique for simultaneous dual-colour tracking of proteins on supported lipid bilayers and on live cells. We show that it out-performs current state-of-the-art labelling while still allowing for convenient one-to-one targeting of molecules via standard tagging approaches. We demonstrate its applicability for the study of molecular interactions at 2D interfaces by quantifying 2D-*KD* constants for two FKBP dimerisers. We expect DNA-PAINT-SPT to work with a wide range of other common tagging approaches in addition to those tested in this study. DNA-PAINT-SPT is versatile and easy to implement by the single-molecule community, given that it uses standard tags and that a wide range of DNA modifications and fluorophores are commercially available. In the future, further improvements of organic dyes will directly benefit DNA-PAINT-SPT and allow for tracking with even higher spatiotemporal resolution.

## ACKNOWLEDGEMENTS

We thank Elia Escoffier for testing data analysis code, Elisa Teunissen for maintaining cell lines and performing protein purification. We acknowledge support from AMOLF technical engineering departments.

## AUTHOR CONTRIBUTIONS

C. Niederauer conceived and performed live cell and *in vitro* experiments, analyzed and interpreted data and wrote the manuscript. C. Nguyen performed *in vitro* experiments. M. Wang-Henderson performed initial *in vitro* experiments. C. Niederauer and M. Wang-Henderson wrote analysis code. J. Stein and F. Stehr performed initial experiments. S. Strauss performed nanobody-DNA coupling and reviewed the manuscript. A. Cumberworth derived analytical homodimerisation model. J. Stein, A. Cumberworth, R. Jungmann and P. Schwille interpreted data and reviewed the manuscript. K.A. Ganzinger conceived and supervised the study, interpreted data and wrote the manuscript.

## DATA AND CODE AVAILABILITY

The raw data that support the findings of this study are available upon reasonable request. Source data and Python code written for the analysis of the data will be available at a public repository.

## Methods

### Materials

Chemicals and materials used were HEPES (H3375, Sigma-Aldrich), sodium chloride (310166, Sigma-Aldrich), magnesium chloride (M8266, Sigma-Aldrich), phosphate-buffered saline tablets (P4417, Sigma-Aldrich), pyranose oxidase (P4234, Sigma-Aldrich), catalase (C40, Sigma-Aldrich), Trolox (238813, Sigma-Aldrich), BSA (A9418, Sigma-Aldrich), Uvasol chloroform (1.02447, Sigma-Aldrich), Uvasol methanol (1.06002, Sigma-Aldrich), sulfuric acid (258105, Sigma-Aldrich), hydrogen peroxide (216763, Sigma-Aldrich), fibronectin (F0895, Sigma-Aldrich), PLL (P4707, Sigma-Aldrich), PLL-PEG-RGD (PLL(20)-g[3.5]-PEG(2)/PEG(3.5)-RGD, SuSoS), His-tagged ICAM-1 (IC1-H52H5, ACROBiosystems), B/B homodimeriser (AP20187, Takara Bio), dibenzocyclooctyne-PEG4-maleimide (760676, Sigma-Aldrich), SNAP/CLIP-tag monoclonal antibody (6F9, Chromotek), single-domain antibody (nanobody) against GFP (clone 1H1, Nanotag Biotechnologies), SNAP-Surface Alexa Fluor 647 (S9136S, New England Biolabs), SNAP-Surface ATTO488 (S9124S, New England Biolabs), Tetraspeck Microspheres 0.2 µm (T7280, ThermoFisher Scientific). SNAPtag-ligand Janelia Fluor dyes (BG-JF549i, BG-JF646, BG-JFX650) were kindly provided by Luke Lavis (Janelia Labs, HHMI).

Lipids used were DOPC (850375, Avanti), DGS-NTA-Ni (790404, Avanti), DSPE-RGD (870295, Avanti), DSPE-PEG-cRGDyk (LP096262-2K, Biopharma PEG) and DOPE-ATTO390 (390-161, ATTO-TEC).

All DNA oligonucleotides were obtained HPLC-purified from Eurofins, except for BG-modified docking strands (Biomers) and azide-modified docking strands (Metabion) used for nanobody-DNA conjugation.

Cell biology media and supplements used were DMEM without Phenol Red (12-917F), DMEM with Phenol Red (12-604F, Lonza), RPMI 1640 with Phenol Red (L0500-500, biowest), RPMI 1640 without Phenol Red (11835-030, gibco), Fluorobrite DMEM (A18967-01, gibco), Pen-Strep (15140122, Sigma-Aldrich), Na-Pyruvate (BE13-115E, Lonza), L-Glutamine (25030-024, gibco), FBS (A3160802, gibco), Ultramem (BE12-743F, Lonza), GeneJuice (70967, Sigma-Aldrich).

### Molecular biology

*pHR-SNAP-CD86-mOrange-FKBP* plasmid was a kind gift from Ricardo A. Fernandes. For live-cell imaging, *pHR-SNAP-CD86-eGFP-FKBP* and *pHR-eGFP-CD86-HaloTag-FKBP* were created using Gibson assembly. For AP20187-induced homodimerisation of FKBP, we introduced a point-mutation (FKBP^f36v^) following previously published protocols ^1,2^ and created *pET30-10His-FKBP*^*f36v*^*-SNAP* using Gibson assembly.

### Cell biology

Jurkat cells were cultured in RPMI with Phenol Red, supplemented with 10% FBS, 1% PenStrep and 1% Na-Pyruvate. HEK cells for lentivirus production were cultured in DMEM with Phenol Red, supplemented with 10% FBS, 1% PenStrep and 1% Glutamax. Phenol-Red free media were used for seeding Jurkat cells into 6-well plates before transfection and labelling.

### Transduction of Jurkat T cells

Jurkat cells were transducted using lentiviral transfection: To this end, HEK293T cells were first transfected with the pHR plasmid of interest, psPax and pMD2G using GeneJuice, following the manufacturer’s protocol. Viral supernatant was collected after 72h and used for Jurkat transduction.

### Nanobody-DNA conjugation

Nanobodies with a single ectopic cysteine at the C-terminus for site-specific conjugation were conjugated similarly to the method described previously ^3^. Unconjugated nanobodies were thawed on ice, then 20-fold molar excess of bifunctional maleimide-Peg4-DBCO linker was added and reacted for 2h on ice. Unreacted linker was removed by buffer exchange using Amicon centrifugal filters (10,000 MWCO). Then, two equivalents of azide-functionalised DNA (5’) were reacted with the DBCO-modified nanobodies overnight at 4 °C. Unconjugated protein and free DNA were removed by anion exchange chromatography using an ÄKTA pure system equipped with a Resource Q 1mL column.

### Recombinant protein expression

10His-FKBP^f36v^-SNAP was expressed in *E*.*coli* (Rosetta strain), purified via its His-tag using an ÄKTA pure system equipped with a HisTrap HP 1mL column, followed by size-exclusion chromatography using a Superdex 200 increase 10/300 GL column.

### Small-unilamellar vesicle generation

Lipids were dissolved in chloroform (to dissolve DSPE-PEG-RGD, 10% methanol were added, and the mixture was sonicated for 30 s in a bath sonicator) and stored in 1.5 mL glass vials with PTFE-lined caps at −20 °C. Lipid mixes were prepared from the stock solutions depending on the required bilayer composition (reconstitution experiments: 98.5% DOPC, 1% DGS-NTA-Ni, 0.5% DOPE-ATTO390; live-cell experiments: 89.5% DOPC, 10% DSPE-PEG-RGD, 0.5% DOPE-ATTO390) and 1 mL were transferred to a 50 mL round bottom flask. By gentle swirling and nitrogen flow, the lipid was dried into a thin film onto the flask walls. Once dried, trace amounts of chloroform were removed by desiccating the flask for at least 2h protected from light.

Afterwards, the dried lipid film was rehydrated in HBS (HEPES 40 mM, pH 7.6, NaCl 140 mM) at a concentration of 2 mg/mL, aliquoted and stored at −20 °C until further use. On the day of the experiment, aliquots were thawed and sonicated for 30 min in a bath sonicator to produce small-unilamellar vesicles (SUVs). SUVs were diluted towards 0.1 mg/mL and used on the same day.

### Preparation of imaging chambers

Coverslips with the dimensions 25 × 75 mm, 1.5H (10812, Ibidi) and 22 × 22 mm, 1.5H (631-0851, VWR) were piranha cleaned using H2SO4 and H2O2 in a 3:1 ratio. After 1h, they were thoroughly rinsed with milliQ water and dried using nitrogen flow. Slides were then air-plasma cleaned for 10 minutes (Harrick Plasma Cleaner PDC-002-HPCE). Chambers were created either by adhering ibidi sticky-slide 8-well or 18-well chambers (80808 or 81818, ibidi) onto pre-treated glass coverslips, or glueing 0.5 mL Eppendorf tubes with the conical part cut off onto coverslips, using UV-curable optical adhesive (NOA68, Thorlabs) and a 36 W UV nail dryer (B00R4M0TI0, Nailstar).

Immediately after plasma cleaning and chamber assembly, 50 µL of HBS were added to each chamber. Then, 50 µL of the respective SUV solution (0.1 mg/mL) were added and the samples were transferred to a moisturised box for 30 minutes. Chambers were washed with 2 mL HBS and bilayers were blocked using BSA (1% (w/v) in HBS) for 10 minutes. Chambers were washed again with 2 mL HBS and stored in a dark moisturised box until further use on the same day. For the screening of surface passivation methods, fibronectin (100 µg/mL, 1h incubation), PLL-PEG-RGD (0.8 mg/mL, 2h incubation), PLL (100 µg/µl, 1h incubation) and His-tagged ICAM-1 (20 nM, 1h incubation on SLB containing 1% DGS-NTA-Ni) were used.

### Protein labelling and reconstitution in SLBs

FKBP^f36v^was thawed on ice and centrifuged for 1h at 4 °C at 16100 x *g*. For dual-colour experiments, supernatant (2.1 µM) was divided into two aliquots and incubated separately with the respective docking strand (3.6 µM) for 1h at room temperature. Protein-DNA was diluted in HBS with 0.1% BSA and 10 mM MgCl2 and incubated on the prepared lipid bilayers for 1h in a dark moisturised box at room temperature. Chambers were washed with 2 mL HBS and imager strand solution containing 40 nM of the respective imager strands, 5 mM MgCl2, 3.7U/mL pyranose oxidase, 200U/mL catalase, 0.8% glucose, 0.1% (w/v) BSA and 2 mM Trolox-Trolox-quinone (Trolox to Trolox-quinone ratio 10% to 20%, determined via NanoDrop 2000 absorption ^4^), were added.

### Cell labelling and preparation for microscopy

Cells were incubated with BG-DNA docking strands (1 nM to 10 nM), nanobody-docking strands (5 pM to 10 pM) or BG-fluorophores (1 nM to 10 nM) for 30 minutes at 37 °C, 5% CO2. Then, cells were washed three times by centrifugation (5 min, 100 x *g*) and re-suspending in PBS. The final resuspension step was performed in PBS with 5 mM MgCl2 and 0.1% (w/v) BSA and, if applicable, the respective imager strand concentration: DNA-PAINT-SPT experiments were performed with 40 nM imager strands; for DNA single-dye experiments, 100 pM of complementary fluorescently labelled strands were added and washed out after 10 min of incubation. Cells with labelling densities of around 0.1 µm^−2^ were used for analysis.

### TIRF Microscopy

Fluorescence imaging was performed on a custom-built microscope in an objective-type TIRF configuration with an oil-immersion objective (CFI Apochromat TIRF 60x, NA 1.49, Nikon) and a three-colour detection scheme. The optical path and a detailed list of components can be found on https://ganzingerlab.github.io/K2TIRF/K2TIRF/index.html. A pre-assembled laser combiner was used to provide four excitation wavelengths (C-FLEX laser combiner, Hübner Photonics; 405 nm 140 mW, 488 nm 200 mW, 561 nm 220 mW, 638 nm 195 mW). The excitation beam was delivered to the optical bench via a single-mode polarisation-maintaining fiber (kineFLEX-HPV-P-3-S-405..640-0.7-0.7-P0, Qioptiq). The laser light was re-collimated after the fiber using an achromatic doublet lens (*f* =50 mm) and directed through an achromatic quarter-waveplate to ensure circular polarisation. The laser beam was spectrally cleaned using a quad-line bandpass (ZET405/488/561/640xv2, Chroma) and then transformed into a collimated flat-top profile using a refractive beam shaping device (piShaper 6_6_VIS, AdlOptica) ^5^. The laser beam diameter was magnified by a factor of 2.5 using a telescope assembly (*f*_1_ =100 mm, *f*_2_ =−40 mm).

The laser light was focused onto the objective’s back focal plane using an achromatic doublet lens (*f* =250 mm). A stage (KMTS25E/M Motorised Translation Stage, Thorlabs) translated the excitation beam off-axis to switch between widefield, HILO or TIRF imaging. A short penetration depth of the evanescent field was ensured by translating the excitation beam in the back focal plane to the maximum possible value without clipping the beam. The angle of incidence was determined using a sample with fluorescent dye in solution (1 µM Cy3B-conjugated DNA), as previously described ^6^: a circular aperture was placed in the beam path and the lateral displacement of the illuminated circle was measured upon translating the sample along the *z*-axis.

The excitation beam was directed towards the objective by a four-colour notch dichroic mirror (ZT405/488/561/640rpc-UF2, Chroma). Fluorescence emission passing through this dichroic mirror was spectrally filtered with a quad-line notch filter (ZET405/488/561/640mv2, Chroma) and was directed through a tube lens (TTL200-A, Thorlabs). The dichroic mirror, the objective, the tube lens and the quad-band notch filter were all placed in a CNC-milled cube based on the miCube design ^7^. This block also supported a piezo stick-slip stage (SLS-5252, Smaract) to move the sample in *x*-*y*-*z*. The tube lens formed an image outside of the cube, where a custom-built slit aperture was used to crop the image horizontally to enable simultaneous three-colour imaging. In a 4f-system (*f* =300 mm), the fluorescence emission was split spectrally using two dichroic mirrors (ZT561rdc and ZT640rdc, Chroma), filtered using respective bandpass filters (525/30 Brightline, Semrock; ET595/50m, Chroma; 680/42 BrightLine, Semrock) and imaged on a sCMOS camera (primeBSI, Teledyne Photometrics). Individual lenses (*f* =300 mm) on the imaging side of the 4f system ensured matching focal planes for all three channels. The imaging setup resulted in an effective pixel size of 108 nm. Focus stabilisation was achieved using a system based on the pgFocus device ^8^. An infrared laser (CPS808S, Thorlabs) was attenuated using a neutral density filter (NE13A-B, Thorlabs), coupled into the excitation path using a long pass dichroic mirror (ZT775sp2-2p-UF1, Chroma) and focused onto the back focal plane of the objective using a *f* =500 mm lens. Using a manual micrometer stage, the infrared laser was brought into total-internal reflection. The back reflection was filtered through a bandpass filter (FB800-40, Thorlabs) and focused (*f* =200 mm) onto a linescan sensor (TSL1401, Parallax). A feedback loop with the piezo-driven stage moving the sample allowed for focus stabilisation throughout extended measurement durations. The setup was controlled using C++ software developed by Marko Seynen (AMOLF, Software Engineering Department).

### Imaging conditions

Fluorescence microscopy data was recorded at room temperature (22 ± 1 °C) with our custom-built setup operating in three-colour simultaneous imaging mode. To this end, the sCMOS camera readout was cropped to 682 × 2048 pixels, providing a 682 × 682 readout for each channel. The camera was operated at 32.4 ms (*in vitro* experiments) or 72.4 ms (live-cell experiments) exposure times, with frame rates of 25 or 12.5 per second, respectively. The read-out rate was set to 100 MHz and the dynamic range to 12 bit. We performed experiments at laser excitation powers in the range of 4 mW to 40 mW (measured just before the back focal plane of the objective), which translate to irradiances of 15 W/cm^2^ to 150 W/cm^2^ in our setup. Laser powers were kept as low as possible while still allowing for robust localisation above background levels, resulting in a typical localisation precision between 20 nm and 30 nm for all labels. The angle of incidence was 76°, resulting in an evanescent field penetration depth in the range of 68 nm (561 nm laser excitation) to 78 nm (638 nm laser excitation).

### Data analysis

Raw data were localised using *picasso* ^9^. Optical distortions were determined using a calibration slide with 200 nm Tetraspeck multi-colour fluorescently-labelled beads, and corrected using Zernike polynomial gradients ^10^. Trajectories were reconstructed from localisations using *trackpy* (http:////soft-matter.github.io and *Swift 0*.*4*.*3* ^11,12^. Trajectories with durations of less than 10 frames or diffusion constants smaller than 0.01 µm^2^/s were rejected. For analysis of live cell data, regions of interest were selected in *Fiji* ^13^ based on the cell outline of a maximum intensity projection of the underlying video or based on the GFP signal of the cell. Colocalisation was determined by calculating intramolecular distances between localised molecules in the different colour channels. Pairs with distances below 300 nm were marked as interaction candidates. Co-diffusing pairs that colocalise for a duration of at least 10 frames were marked as interacting, and gaps of up to 6 frames were closed. Integration of the different analysis packages, any further analysis and visualisation of data was performed with custom-written *Python*-code soon available at https://github.com/GanzingerLab.

## Supplementary Information

### A. DNA sequences

DNA-sequences are in 5’-3’ notation.

- DNA-PAINT-SPT sequences Imager strand A, Cy3B-conjugated: 4x(GA) -Cy3B
  GAGAGAGA -Cy3B
  Imager strand B, ATTO643-conjugated: 4x(GT) -ATTO643
  GTGTGTGT -ATTO643
  Docking strand A: BG-or azide-conjugated: 37.5x(TC)
  BG/azide -TCTCTCTCTCTCTCTCTCTCTCTCTCTCTCTCTCTCTCTCTCTCTCTCTCTCTCTCTCTCTCTCTCTCTCTCTCT
  Docking strand B: BG-or azide-conjugated: 37.5x(AC)
  BG/azide -ACACACACACACACACACACACACACACACACACACACACACACACACACACACACACACACACACACACACACA
- Single-dye DNA sequences We used a single-dye control designed to be of similar size and mass compared to a 50% imager strand-occupied DNA-PAINT-SPT docking strand. The single-dye docking-imager strand pair consists of a 75 nucleotide docking strand and a 35 nucleotide fully complementary imager strand. At our experimental conditions (ionic strength, temperature, timescale), the imager strand is considered to be irreversibly bound to the docking strand. Imager strand A:
  ATAATAAGTAATCTACAACAATCGGGTGGGTCAGC -Cy3B
  Imager strand B:
  TAATGAAATGGGAACTAACTCTCGGAAACCTTTAT -ATTO643
  Docking strand A:
  BG -TTTTTTTTTTATAAAGGTTTCCGAGAGTTAGTTCCCATTTCATTATTTTTTTTTTTTTTTTTTTTTTTTTTTTTT
  Docking strand B:
  BG -TTTTTTTTTTGCTGACCCACCCGATTGTTGTAGATTACTTATTATTTTTTTTTTTTTTTTTTTTTTTTTTTTTTT

### B. Analytical derivation of dimer concentration for ligand-induced homodimerization

We briefly outline the analytical derivation of the dimer concentration for ligand-induced homodimerization. For a detailed discussion refer to Binder et al., 2021^1^.

We use 2D densities in 1 µm^−2^ and convert molar 3D concentrations into 1 µm^−3^, where

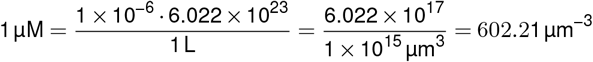

Consider a system with ligands in solution with concentration *c*_*L*_ and protein monomers embedded in a lipid bilayer with surface concentration Γ_*M*_. In this system, the ligands can bind up to two monomers simultaneously. We consider both the equilibrium of a monomer binding to a free ligand (Eq. 1) and the equilibrium of a monomer binding to an already bound ligand (Eq. 2):

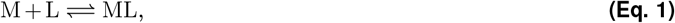

And

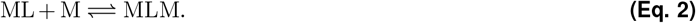

The dissociation constant *K*_*B*_ for a monomer bound to a single ligand, ML, is defined as

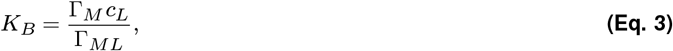

where Γ_*ML*_ is the surface concentration of ML. The dissociation constant *K*_*X*_ for the ligand-homodimer complex, MLM, is defined as

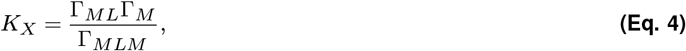

where Γ_*M LM*_ is the concentration of MLM.

To derive an expression for Γ_*M LM*_ that does not include Γ_*M*_ or Γ_*ML*_, we need the equation for the mass balance of the protein monomer,

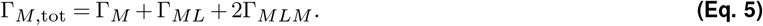

Combining (Eq. 3) to (Eq. 5) and rearranging, we obtain

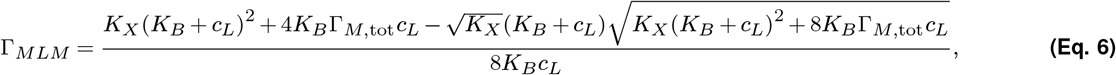

| Surface | Imager strand | Localisations (background) [$\mu\text{m}^{-2}$] | Localisations (cells) [$\mu\text{m}^{-2}$] | Ratio |
| --- | --- | --- | --- | --- |
| BSA | ATTO643 | 0.67 | 0.78 | 1.16 |
|  | Cy3B | 0.25 | 0.37 | 1.44 |
| Fibronectin | ATTO643 | 0.30 | 0.35 | 1.17 |
|  | Cy3B | 0.17 | 0.22 | 1.33 |
| PLL | ATTO643 | 0.17 | 0.30 | 1.81 |
|  | Cy3B | 0.04 | 0.15 | 4.07 |
| PLL-PEG-RGD | ATTO643 | 0.05 | 0.34 | 6.48 |
|  | Cy3B | 0.06 | 0.37 | 6.45 |
| DSPE-RGD | ATTO643 | 0.01 | 0.12 | 9.21 |
|  | Cy3B | 0.01 | 0.08 | 11.08 |
| ICAM | ATTO643 | 0.01 | 0.11 | 15.04 |
|  | Cy3B | 0.01 | 0.10 | 12.34 |
| DSPE-PEG-RGD | ATTO643 | 0.01 | 0.15 | 13.05 |
|  | Cy3B | 0.01 | 0.11 | 16.53 |

where we have discarded the second solution in which the second term (containing the square root) is positive, as it leads to negative values of Γ_*M*_ and Γ_*ML*_.

The density of reconstituted proteins is measured by counting receptors before adding the dimer-inducing ligand, since we can not easily distinguish between a monomer and a same-colour homodimer. One hour after adding ligands, the number of dimers is determined and the fraction of dimerised molecules is fit by (Eq. 6) with free parameters *K*_*X*_, *K*_*B*_ and a correction factor accounting for unlabelled molecules.

### C. Surface passivation

The high concentration of fluorescently labelled imager strands in solution poses two challenges: imager strands bind to the surface via unspecific interactions with the surface or cell debris, and they contribute to a diffuse background, decreasing the signal-to-background ratio. We screened various surface passivation methods (see Fig. S7) and found lipid bilayers to provide the best passivation efficiency. Cell attachment is ensured by functionalising the bilayers with cell adhesion promoting molecules (RGD, PEG-RGD functionalised lipids or His-tagged ICAM-1 reconstituted on nickelated lipids). Bilayers containing PEG-RGD functionalised lipids or RGD-functionalised lipids had comparable passivation efficiencies, but PEG-RGD functionalised lipids performed better in terms of cell attachment. We also used PLL-PEG-RGD successfully for single-colour experiments with Cy3B. When used with ATTO643-labelled imager strands, significant binding of imager strands to the surface was observed, which could at least partially be prevented by adding Trolox/Trolox-quinone as a reductant and oxidant system ^2^.

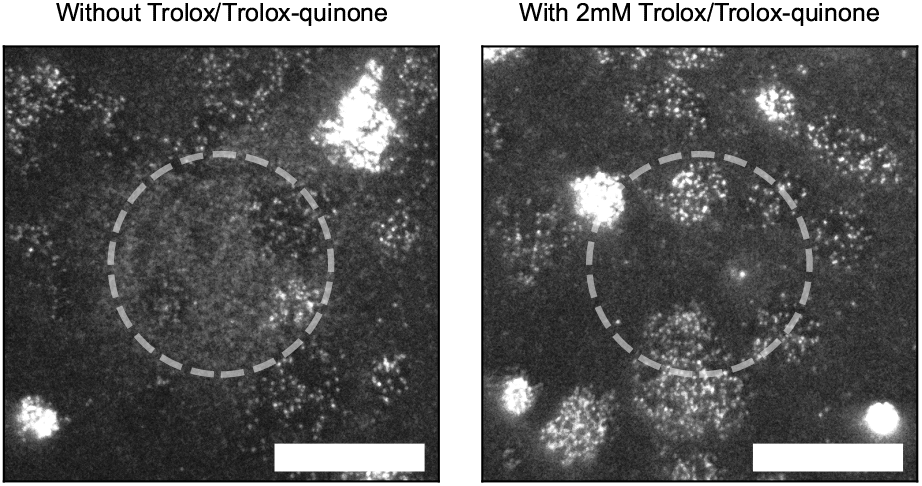

### ATTO643 photoreaction on PLL-PEG-RGD with and without Trolox

Cells labelled with DNA-PAINT and 40 nM ATTO643-labelled imager strands. First frame after illuminating for 1 min in the center circular region, with 2 mM Trolox/Trolox-quinone (left), or without Trolox/Trolox-quinone in the imaging buffer (right).

Scale bar: 10 µm.

## Supplementary Figures

**Figure S1.**
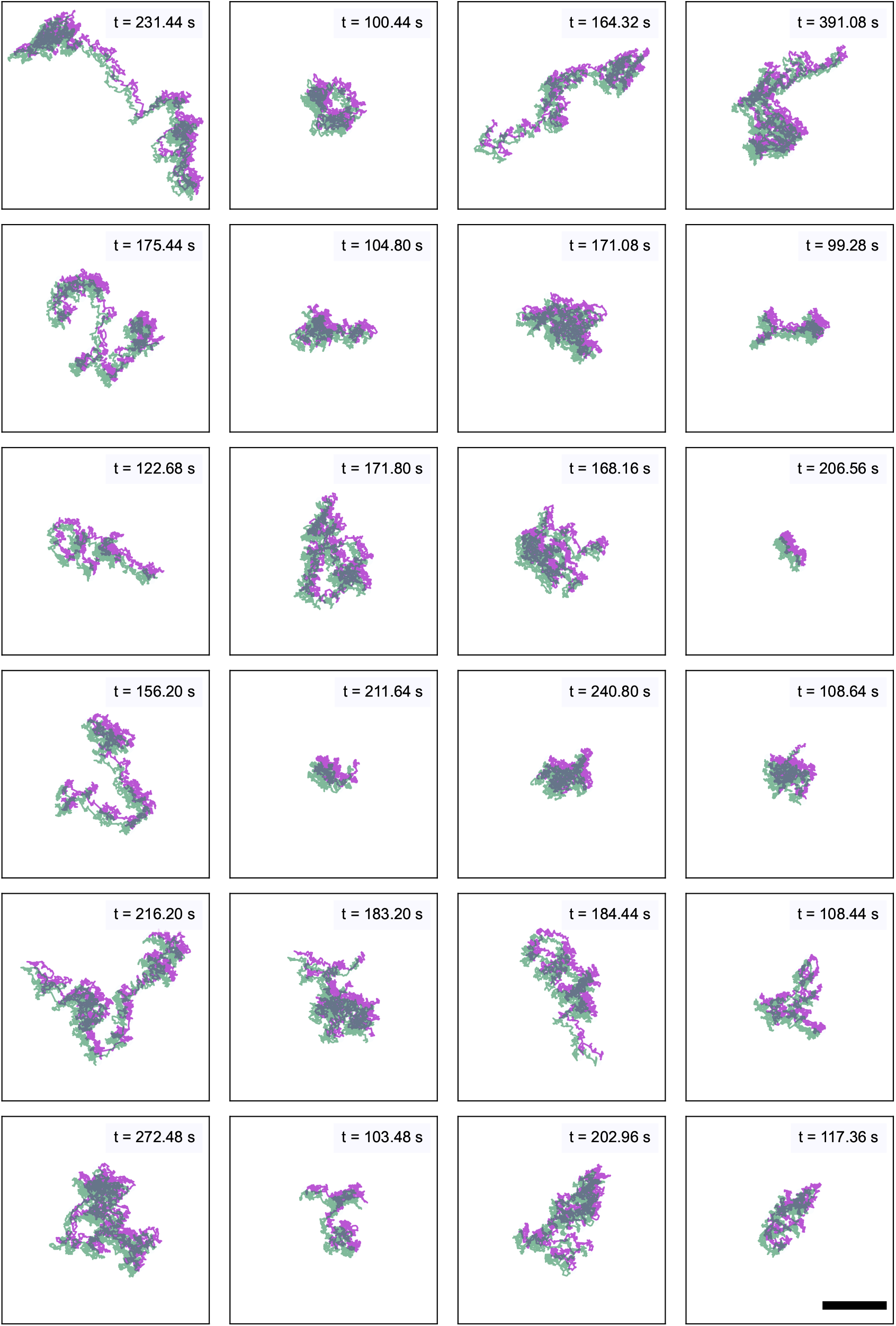
Ligand-induced FKBP dimers detected with dual-colour DNA-PAINT-SPT. The longest trajectories (30th-percentile) of co-diffusing AP20187-induced FKBP dimers, labelled with dual-colour DNA-PAINT and recorded during a 15 minute TIRFM measurement. For displaying purposes, tracks were moved in opposite *x* and *y* directions by 1 µm. Scale bar: 10 µm.

**Figure S2.**
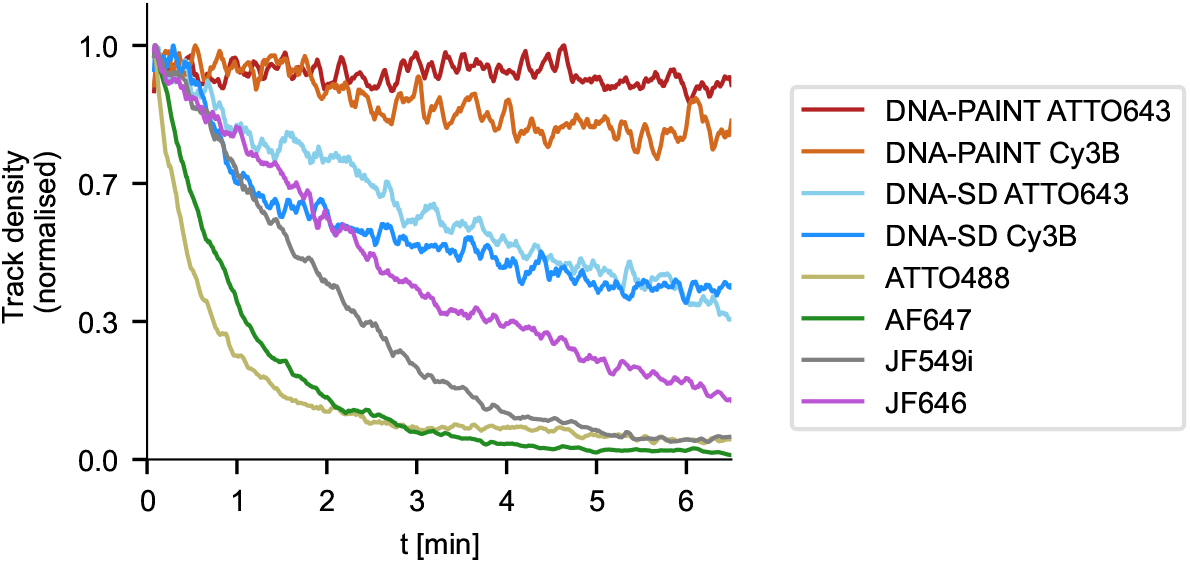
Photostability of different labelling methods and fluorophores in *in vitro* experiments. Number of trajectories per frame of reconstituted FKBP proteins labelled with DNA-PAINT (ATTO643-imager strands in red, *n*_samples_ = 3; Cy3B-imager strands in orange, *n*_samples_ = 3), single-dye DNA (ATTO643-fluorophore in light blue, *n*_samples_ = 4; Cy3B-fluorophore in dark blue, *n*_samples_ = 4) or single BG-conjugated fluorophores (ATTO488 in yellow, *n*_samples_ = 2; Alex-aFluor647 in green, *n*_samples_ = 2; JF549i in gray, *n*_samples_ = 5; JF646 in magenta, *n*_samples_ = 5).

**Figure S3.**
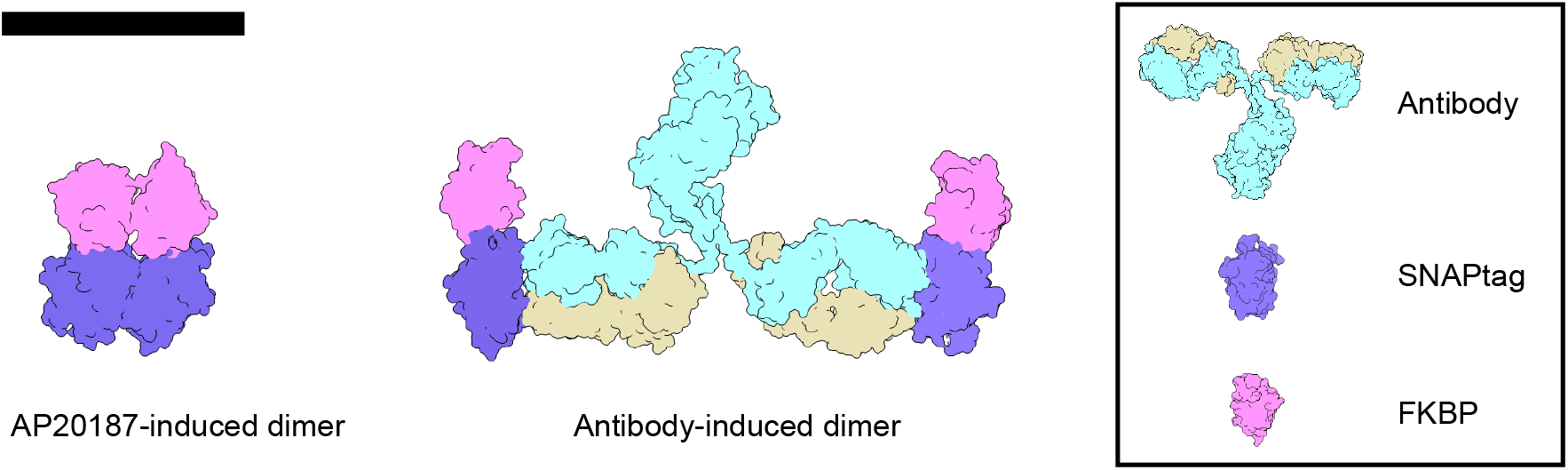
Dimerisation geometry for ligand-and antibody-induced dimerisation. Size comparison of FKBP-SNAPtag complexes dimerised via the ligand AP20187 and an anti-SNAPtag antibody. Protein structures from PDB (SNAPtag: 3KZY, FKBP^f36v^: 1BL4, Antibody: 1IGT). Scale bar: 10 nm.

**Figure S4.**
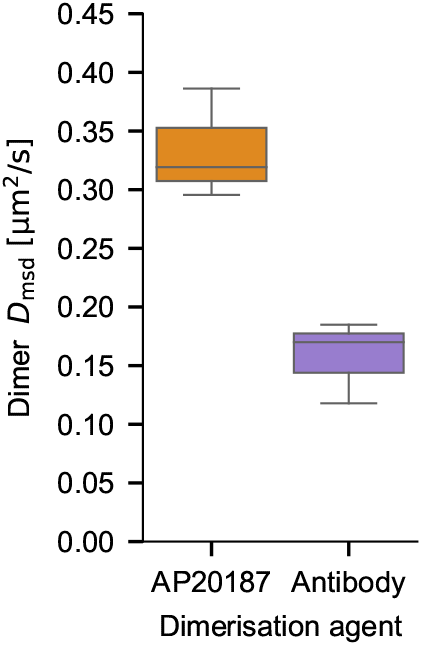
Diffusion constants of dimers for AP20187-and antibody-induced dimerisation. Diffusion constants derived from mean-square displacement of co-diffusing trajectories show a two-fold reduction for antibody-induced dimers (*D*_msd, AP20187_ = 0.33 *±* 0.05 µm^2^ /s, *D*_msd, Antibody_ = 0.16 *±* 0.04 µm^2^ /s). This effect is likely due to the bigger size and mass of the antibody-induced dimer, and potentially also by the increased coupling of thermal energy to the rotational degree of freedom, as the antibody-induced dimer has a higher moment of rotational inertia compared to the compact ligand-induced dimer. Data from three samples for each condition.

**Figure S5.**
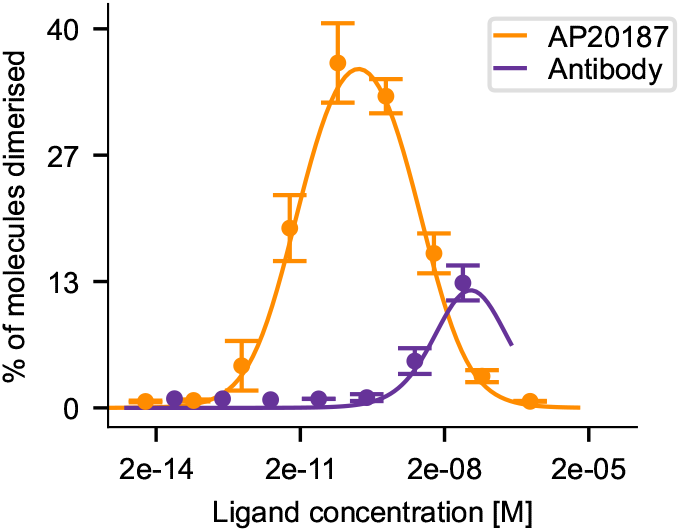
Dimerisation curve of ligand-titration experiments measured using DNA-PAINT-SPT. Fraction of dimerised molecules as detected using DNA-PAINT-SPT during ligand-titration experiments. AP20187 or anti-SNAPtag antibody were used to induce dimerisation. Labelled fractions according to the fit were 60 *±* 17 %. Error bars denote mean *±* standard deviation of data collected from three field of views of each sample of a titration series for each dimerisation agent.

**Figure S6.**
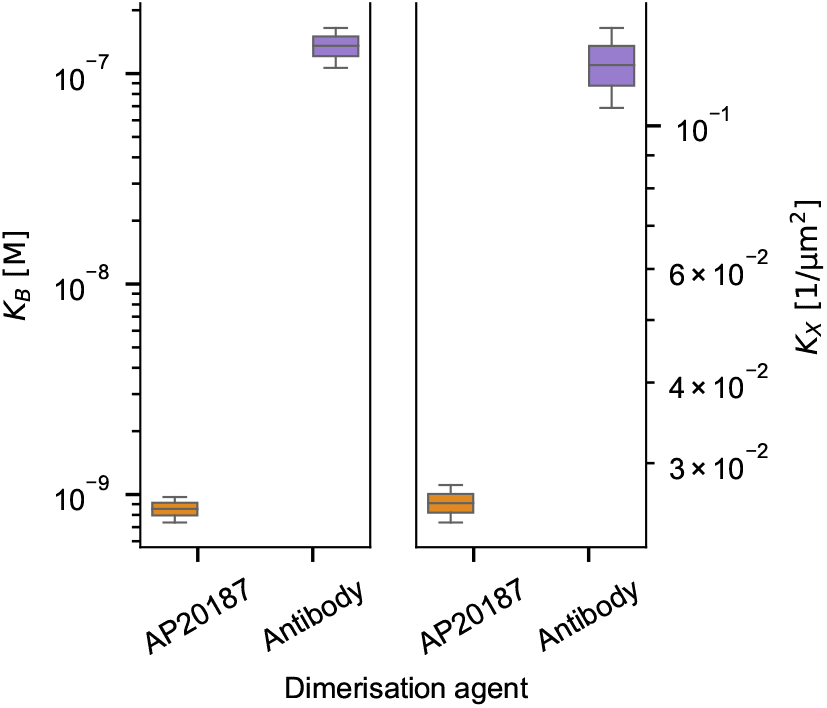
2D dissociation constants of different dimerisation agents measured using DNA-PAINT-SPT. Dissociation constants *K_X_* and *K_B_* of AP20187 and anti-SNAPtag antibody interaction with FKBP^f36v^ as determined from fitting the fraction of dimerised molecules. Labelled fractions according to the fits were 67 *±* 16 %. Boxes, line and whiskers show, respectively, 25–75 quartiles, median, and minimum and maximum values of dissociation constants. The data used for fitting was collected from three field of views of each sample of a titration series, prepared in duplicates for each dimerisation agent.

**Figure S7.**
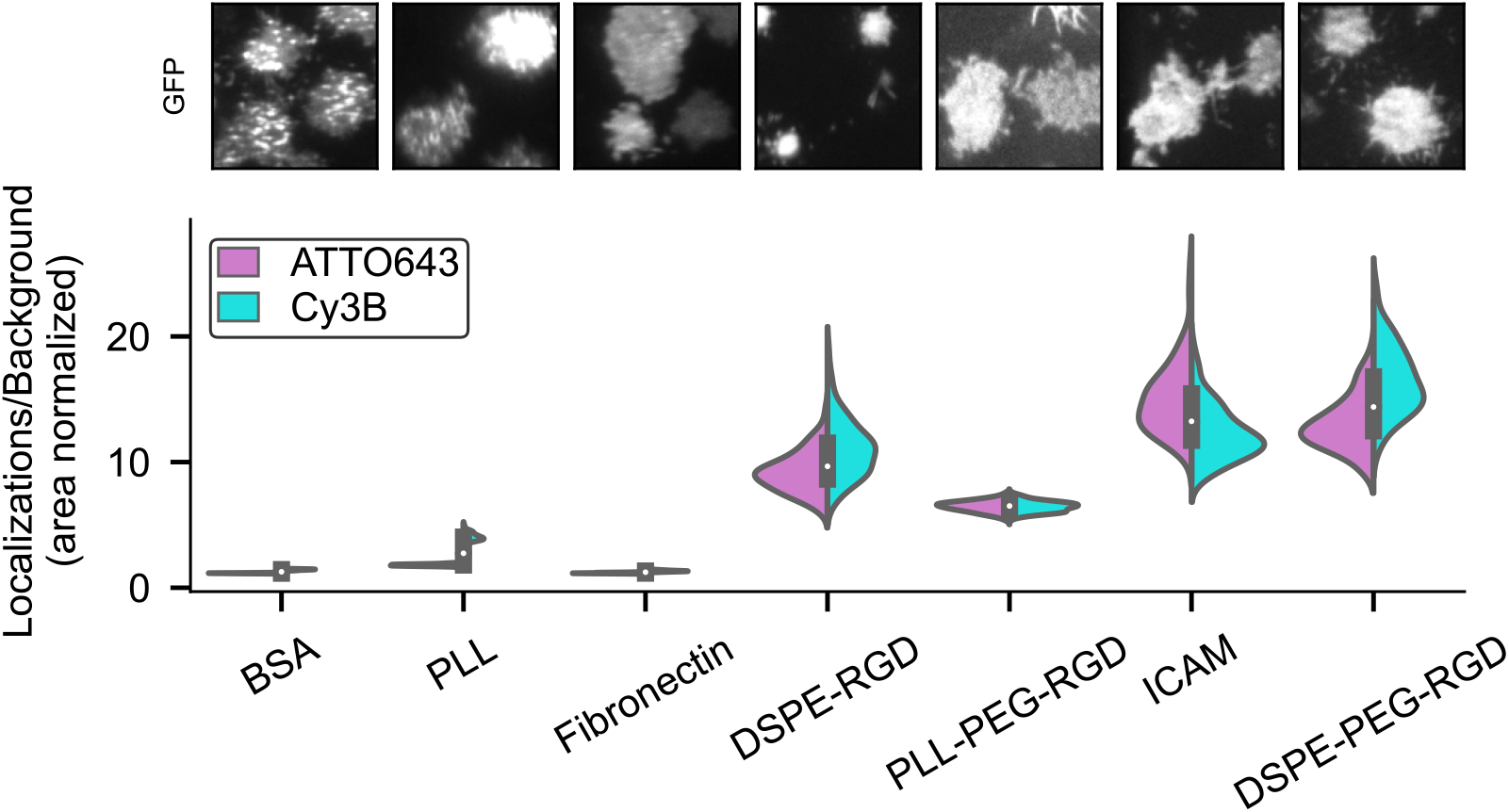
Screening of passivation methods. Ratio of localisations detected on DNA-PAINT labelled cells with 40 nM imager strands (Cy3B-or ATTO643-conjugated) per area versus localisations detected outside of cells per area, for different surfaces. Upper row shows GFP signal of adhered cells (field of view: 20 µm). See **??** for video version with single-molecule channels. Data and panels from one representative field of view for each condition.

**Figure S8.**
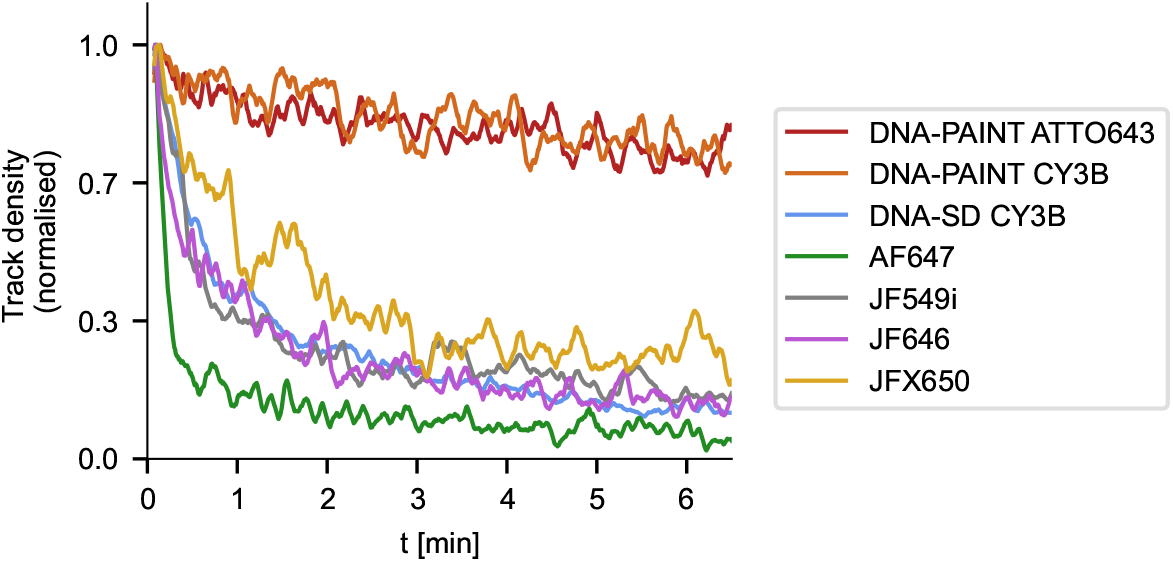
Trajectory density over time for DNA-PAINT labels and single-dye controls. Number of trajectories per frame on individual cells labelled with DNA-PAINT (ATTO643-imager strands in red, *n*_cells_ = 25; Cy3B-imager strands in orange, *n*_cells_ = 25), single-dye DNA (Cy3B-fluorophore in blue, *n*_cells_ = 32) or single BG-conjugated fluorophores (AlexaFluor647 in green, *n*_cells_ = 7; JF549i in gray, *n*_cells_ = 15; JF646 in magenta, *n*_cells_ = 6; JFX650 in yellow, *n* cells = 6).

**Figure S9.**
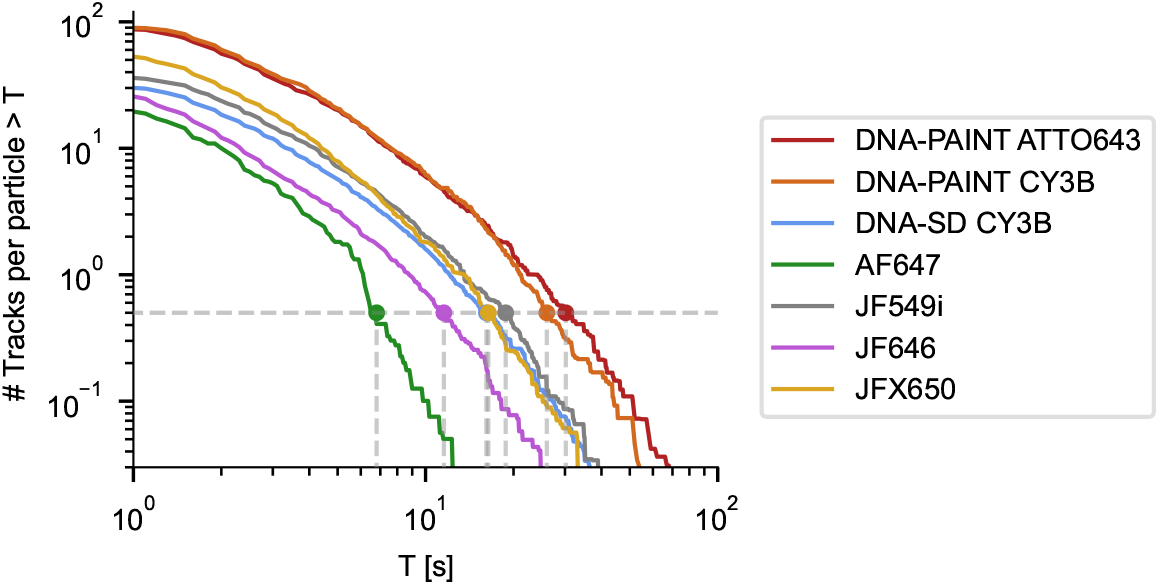
Tracks per particle over time for DNA-PAINT labels and single-dye controls. Single-molecule trajectories with a duration longer than T, normalised to the initial number of trajectories per individual cell. Membrane proteins are labelled with DNA-PAINT (ATTO643-imager strands in red, *n*_cells_ = 25; Cy3B-imager strands in orange, *n*_cells_ = 25), single-dye DNA (Cy3B-fluorophore in blue, *n*_cells_ = 32) or single BG-conjugated fluorophores (AlexaFluor647 in green, *n*_cells_ = 7; JF549i in gray, *n*_cells_ = 15; JF646 in magenta, *n*_cells_ = 6; JFX650 in yellow, *n*_cells_ = 6).

**Figure S10.**
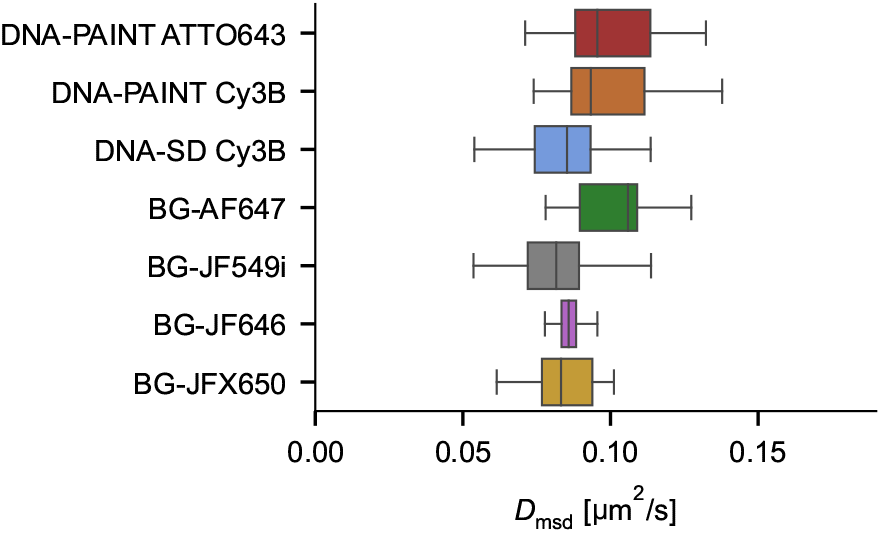
Diffusion constants of membrane proteins labelled with DNA-PAINT and single dye probes. Mean-square displacement derived diffusion constants of membrane proteins on cells labelled with DNA-PAINT (ATTO643-imager strands in red, *n*_cells_ = 25; Cy3B-imager strands in orange, *n*_cells_ = 25), single-dye DNA (Cy3B-fluorophore in blue, *n*_cells_ = 32) or single BG-conjugated fluorophores (AlexaFluor647 in green, *n*_cells_ = 7; JF549i in gray, *n*_cells_ = 15; JF646 in magenta, *n*_cells_ = 6; JFX650 in yellow, *n*_cells_ = 6).

**Figure S11.**
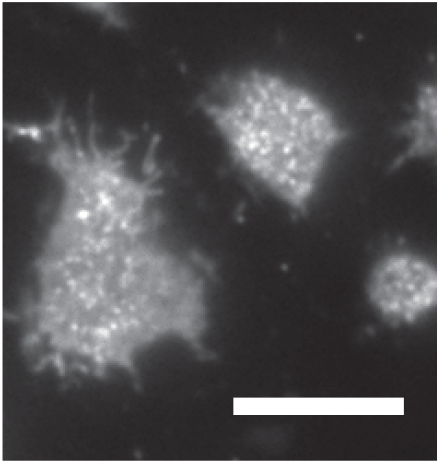
Accessibility of docking strands. Average fluorescence intensity during a 80 s measurement of Jurkat T-cells with densely labelled membrane proteins using DNA-PAINT docking strands and Cy3B-conjugated imager strands, to visualise potential exclusion effects of DNA-labelled proteins from cell-surface contacts.

**Figure S12.**
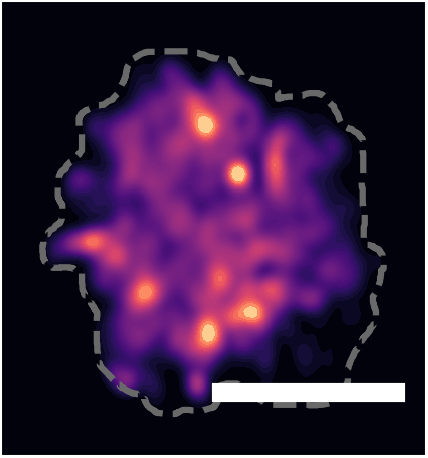
Trajectory density variation across cell surface. Density of single-molecule trajectories across Jurkat T-cell surface during a 400 s measurement for DNA-PAINT docking strand labelled membrane proteins and Cy3B-conjugated imager strands.

## Supplementary Videos

**Figure SV1.**
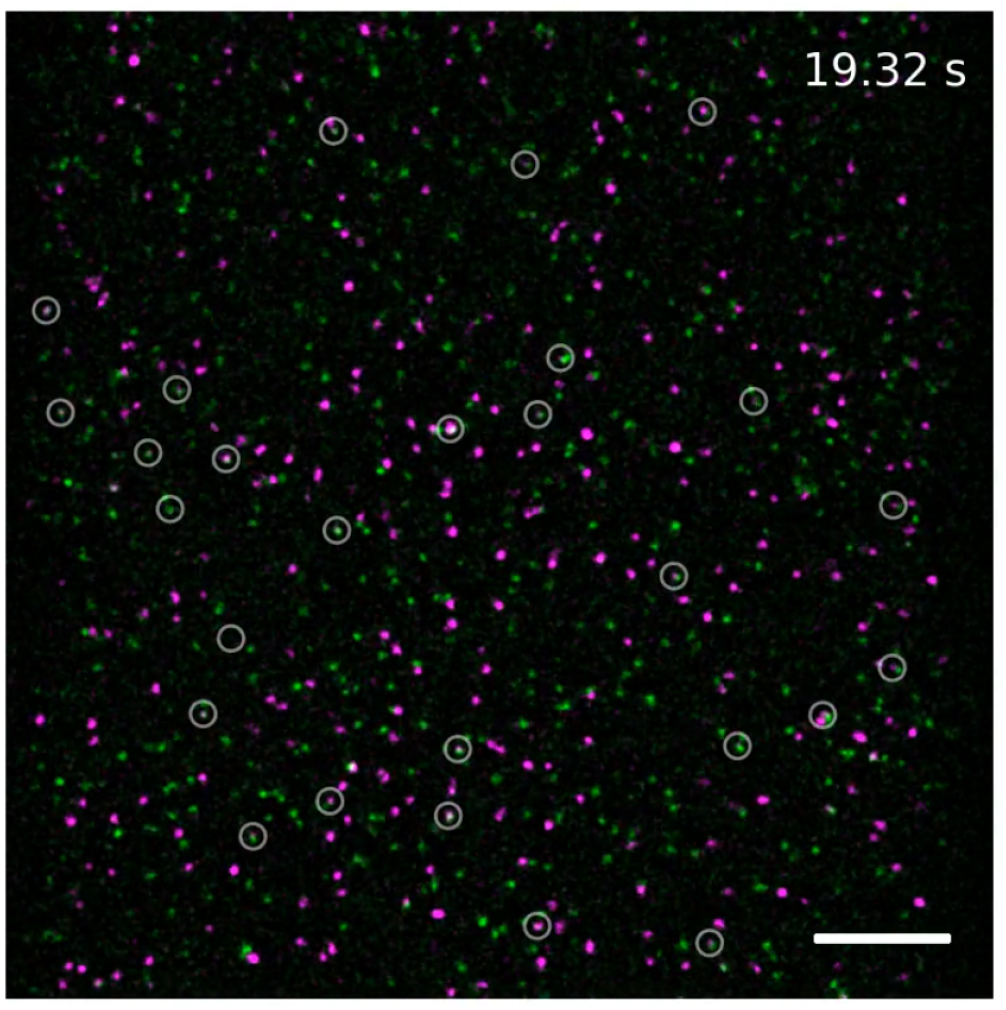
SLB with reconstituted FKBP proteins labelled with dual-colour DNA-PAINT. TIRFM video (40 ms exposure time, replayed at 25 fps) of reconstituted dimerised FKBP proteins labelled with dual-colour DNA-PAINT. Colocalisation events are marked with a circle. Scale bar 10 µm.

**Figure SV2.**
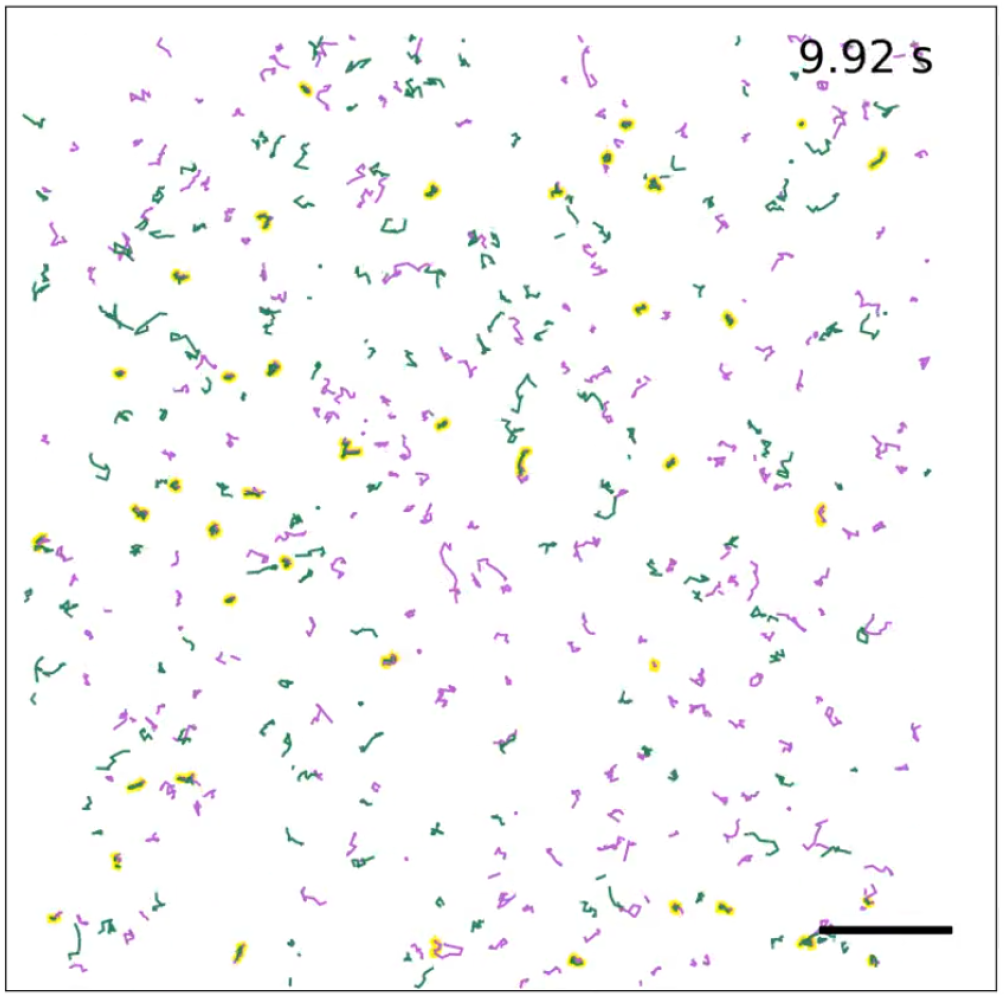
Single-molecule tracking and detection of co-diffusing trajectories. Trajectories of molecules displayed in Fig. SV1 with co-diffusing trajectories shaded in yellow. Scale bar 10 µm.

**Figure SV3.**
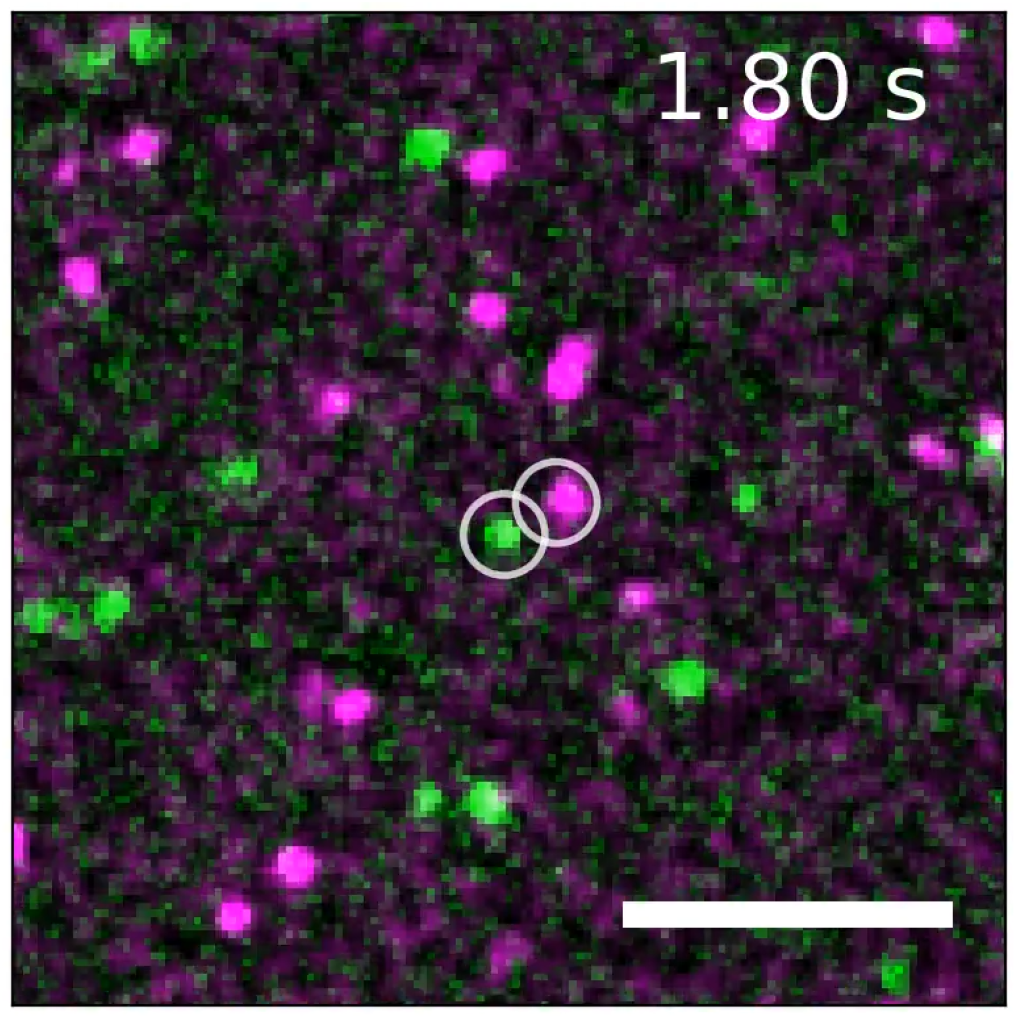
Association and dissociation of a dimer. TIRFM video (40 ms exposure time, replayed at 25 fps) of reconstituted dimerised FKBP proteins labelled with dual-colour DNA-PAINT. Dimerisation is induced by antiSNAP antibodies. Two FKBP monomers associate (white circles, association at *≈* 2.4 s), co-diffuse (blue circle and track) for about 21 s, and dissociate into two monomers (white circles, dissociation at *≈* 23.5 s. Scale bar: 4 µm.

**Figure SV4.**
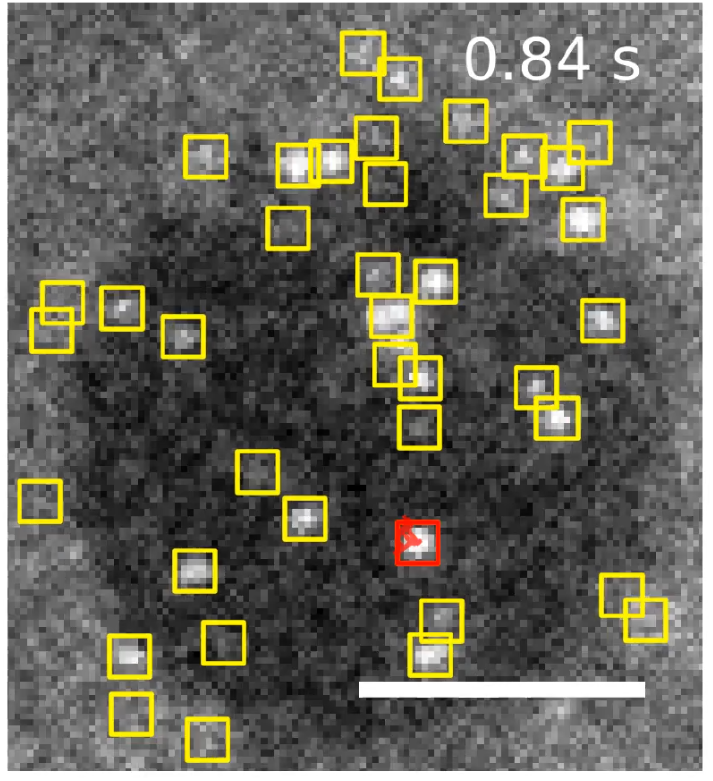
DNA-PAINT labelled membrane proteins diffusing on Jurkat T cell. TIRFM video (80 ms exposure time, replayed at 12 fps) showing single-molecule trajectories of DNA-PAINT labelled membrane proteins expressed on a Jurkat T cell. Localised molecules (yellow boxes) and trajectory of an individual molecule (red box and trajectory). Scale bar: 5 µm.

**Figure SV5.**
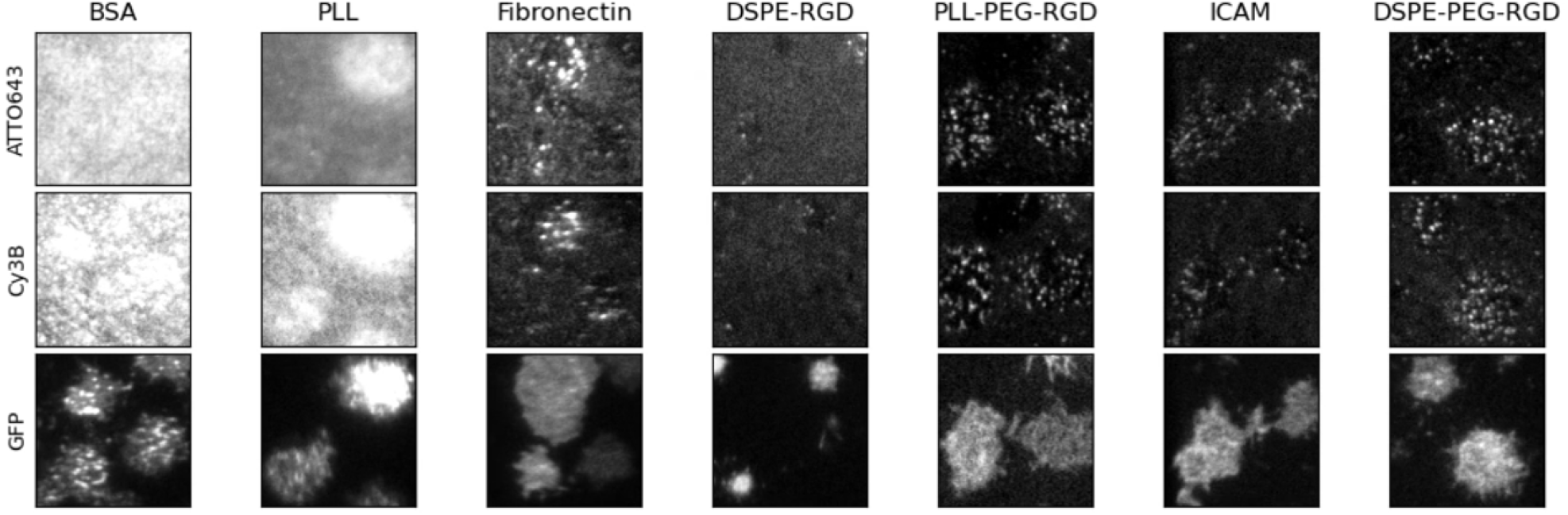
Screening of passivation methods. TIRFM video (80 ms exposure time, replayed at 25 fps) of dual-colour DNA-PAINT-SPT of membrane proteins and GFP signal of adhered cells on different surfaces. Field of view: 20 µm.

**Figure SV6.**
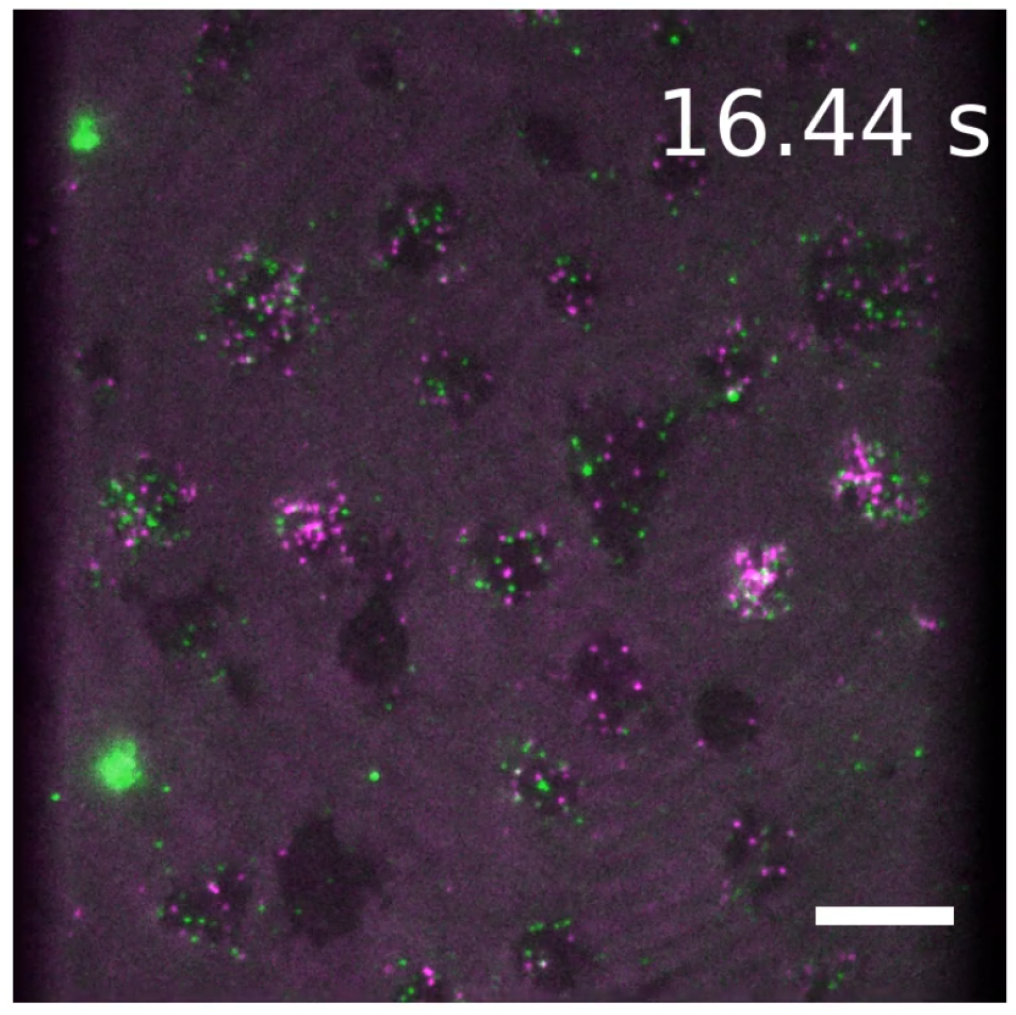
Dual-colour DNA-PAINT labelled membrane proteins diffusing on Jurkat T cell. TIRFM video (80 ms exposure time, replayed at 12 fps) of membrane proteins expressed on a Jurkat T cell and labelled orthogonally with dual-colour DNA-PAINT-SPT. Scale bar: 10 µm.

